# Fast end-to-end learning on protein surfaces

**DOI:** 10.1101/2020.12.28.424589

**Authors:** Freyr Sverrisson, Jean Feydy, Bruno E. Correia, Michael M. Bronstein

**Affiliations:** École Polytechnique Fédérale de Lausanne and Swiss Institute of Bioinformatics; Imperial College London; Imperial College London / Twitter

**Author notes:** equal contribution (Freyr Sverrisson, Jean Feydy).

## Abstract

Proteins’ biological functions are defined by the geometric and chemical structure of their 3D molecular surfaces. Recent works have shown that geometric deep learning can be used on mesh-based representations of proteins to identify potential functional sites, such as binding targets for potential drugs. Unfortunately though, the use of meshes as the underlying representation for protein structure has multiple drawbacks including the need to pre-compute the input features and mesh connectivities. This becomes a bottleneck for many important tasks in protein science.

In this paper, we present a new framework for deep learning on protein structures that addresses these limitations. Among the key advantages of our method are the computation and sampling of the molecular surface on-the-fly from the underlying atomic point cloud and a novel efficient geometric convolutional layer. As a result, we are able to process large collections of proteins in an end-to-end fashion, taking as the sole input the raw 3D coordinates and chemical types of their atoms, eliminating the need for any hand-crafted pre-computed features.

To showcase the performance of our approach, we test it on two tasks in the field of protein structural bioinformatics: the identification of interaction sites and the prediction of protein-protein interactions. On both tasks, we achieve state-of-the-art performance with much faster run times and fewer parameters than previous models. These results will considerably ease the deployment of deep learning methods in protein science and open the door for end-to-end differentiable approaches in protein modeling tasks such as function prediction and design.

## 1. Introduction

Proteins are biomacromolecules central to all living organisms. Their function is a determining factor in health and disease, and being able to predict functional properties of proteins is of the utmost importance to developing novel drug therapies. From a chemical perspective, proteins are polymers composed of a sequence of amino acids (Fig. 1.a). This sequence determines the structural conformation (fold) of the protein, and the structure in turn determines its function. In a folded protein, hydrophobic (water-repelling) residues typically cluster within the core of the protein, while hydrophilic (water-attracting) residues are exposed to water solvent on its surface. The properties of this surface dictate the type and the strength of the interactions that a protein can have with other molecules (Fig. 1.b). Analysing this complex 3D object is therefore a fundamental problem in biology: models for protein structures can be used to understand the possible interactions between a protein and its environment, and consequently predict the functions of these macromolecules in living organisms.

**Figure 1:**
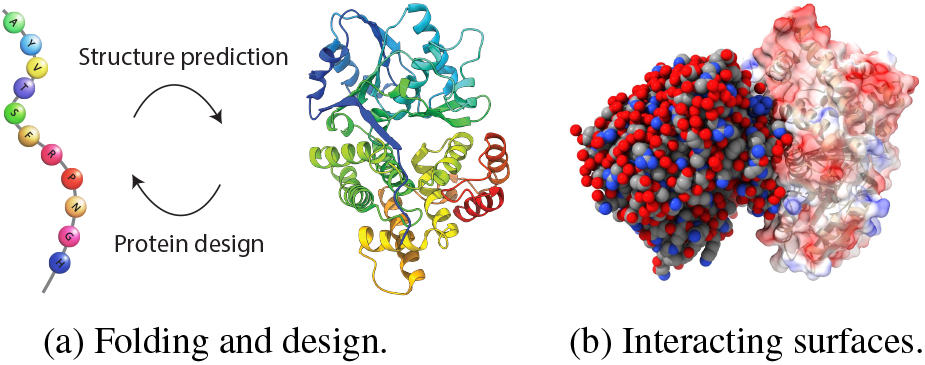
Three major problems in structural biology. (a) Protein design is the inverse problem of structure prediction. (b) Two interacting proteins represented as an atomic point cloud (left) and as a molecular surface (right) that abstracts out the internal fold (shown semi-transparently). Protein surfaces display a number of geometric (e.g. con-cave and convex regions) and chemical (e.g. charges) features. Identifying their binding is a complex problem that can be addressed with geometric deep learning.

Since proteins are predominant drug targets, the study of their interactions with other molecules is a key problem for fundamental biology and the pharmaceutical industry. Classical drugs are small molecules designed to bind to a protein of interest, with a binding site that usually has noticeable ‘pocket-like’ structure. Targets with flat surfaces that exhibit no pockets have long been a challenge for drug developers and are often deemed ‘undruggable’. The possibility of addressing such targets with specifically designed protein molecules (known as biological drugs or ‘biologics’) is a fast emerging field in drug-development holding the promise to provide novel therapeutic strategies for many important diseases (e.g. cancer, viral infections, ect.).

Deep learning methods have increasingly been applied to a broad range of problems in protein science [21], with the particularly notorious success of DeepMind’s AlphaFold to predict 3D protein structure from sequence [37]. Recently, Gainza et al. [20] introduced MaSIF, one of the first conceptual approaches for *geometric deep learning* on protein molecular surfaces allowing to predict their binding. The main limitations of MaSIF stem from its reliance on pre-computed meshes and handcrafted features, as well as significant computational time and memory requirements.

### Main contributions

In this paper, we present dMaSIF (differentiable molecular surface interaction fingerprinting), a new deep learning approach to identify interaction patterns on protein surfaces that addresses the key drawbacks of MaSIF. Our architecture is completely free of any pre-computed features. It operates directly on the large set of atoms that compose the protein, generates a point cloud representation for the protein surface, learns task-specific geometric and chemical features on the surface point cloud and finally applies a new convolutional operator that approximates geodesic coordinates in the tangent space. All these computations are performed on the fly, with a small memory footprint. Notably, we implement all core calculations as reductions of symbolic “distance-like” matrices, supported by the recent KeOps library [19] for PyTorch [30]: the high performance routines of this toolbox allow us to design a method which is fully differentiable and an order of magnitude faster and more memory efficient than MaSIF. This in turn allows us to make predictions on larger collections of protein structures than was previously practical, and opens the door to end-to-end protein optimization and *de novo* protein design using geometric deep learning.

## 2. Related works

### Deep learning in protein science

Proteins can be rep-resented in different ways, the 1D aminoacid sequence being the simplest and most abundant source of data. Recent methods have taken advantage of the wealth of protein sequences available in public databases and shown how unsupervised embeddings borrowed from the field of Natural Language Processing can improve function prediction [2, 8, 36]. Deep learning is also becoming a key component in many pipelines for protein folding (i.e. inferring the 3D structure from the aminoacid sequence) [3, 46, 37, 47]. Many of these pipelines predict pair-wise distances and other geometric relations between different residues and use these as constraints in later structural refinements. Protein design, which can be considered as ‘inverse structure prediction’ (i.e. predict a sequence that will fold into a particular structure), has also benefited from deep learning methods [23]. We refer to [21] for a comprehensive overview.

To model protein interactions, surface-based representations are especially attractive: they automatically abstract the less relevant internal parts of the protein fold, which do not contribute to the interaction. The Molecular Surface Interaction Fingerprinting (MaSIF) [20] method pioneered the use of mesh-based geometric deep learning to predict protein interactions. Its authors showed the application of MaSIF for classifying binding sites for small ligands, discriminating sites of protein-protein interaction in surfaces and predicting protein-protein complexes.

Nevertheless, in spite of its conceptual importance and impressive performance, the MaSIF method has significant drawbacks that limit its practical applications for protein prediction and design. First, it takes as inputs mesh-based representations of a protein surface, that must be generated from the raw atomic point cloud as a preprocessing step. Second, it relies on hand-crafted chemical and geometric features that must also be pre-computed and stored on the hard drive. Third, it uses MoNet [29] mesh convolutions on precomputed geodesic patches, which becomes prohibitively expensive in terms of memory and run time when working with more than a few thousand proteins.

### Deep learning on surfaces and point clouds

Deep learning on non-Euclidean structured data such as meshes, graphs and point clouds, known under the umbrella term *geometric deep learning* [11], has recently become an important tool in computer vision and graphics. Instead of considering geometric shapes as objects in a 3D Euclidean space and applying standard deep learning pipelines (e.g. based on 2D views [44], volumetric [38], space partitioning [35, 42, 39] and implicit representations [14]), geometric deep learning seeks to develop a non-Euclidean analogy of filtering and pooling operations. Boscaini et al. [26] proposed the first geometric CNN-like architecture (Geodesic CNN) based on intrinsic local charting on meshes. Follow-up works improved on these results using patch operators based on anisotropic diffusion (ACNN [10]), Gaussian mixtures (MoNet [29]), splines [17], graph message passing (FeastNet [41]), equivariant filters [31, 15], and primal-dual mesh operators [28]. We refer to [32] for a recent survey.

Point clouds are often used as a native representation of 3D data coming from range sensors, and have recently gained popularity in computer vision in lieu of surface-based representations. First works on deep learning on point clouds were based on deep learning on sets [48] (PointNet and PointNet++ [34]). DGCNN [43] uses graph neural networks [6] on kNN graphs constructed on the fly to capture the local structure of the point cloud. Additional tangent space [39] and volumetric [4] convolution operators were also considered, see a recent survey paper [22].

## 3. Our approach

### Working with protein surfaces

In the following, we describe a new efficient end-to-end architecture for geometric deep learning on protein molecules. The premise of our work is that protein molecular surfaces carry important geometric and chemical information indicative of the way they interact with other molecules. Though we showcase our method on predicting binding properties (arguably, the most important task in structural biology and drug design), it is generic and can be trained on other problems, and in principle, extended to other biomolecules.

Our method works on successive geometric representations of a protein, illustrated in Figure 2. The input is provided as a cloud of atoms {**a**_1_, *…*, **a**_A_} ⊂ ℝ^3^, with chemical types in the list [C, H, O, N, S, Se] encoded as one-hot vectors {**t**_1_, *…*, **t**_A_} ⊂ ℝ^6^. We then represent the surface of the protein as an oriented point cloud {**x**_1_, *…*, **x**_N_} ⊂ ℝ^3^ with unit normals 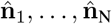 in ℝ^3^. We associate feature vectors **f**_1_, *…*, **f**_N_ to these points and progressively update them by convolution-like operations; the dimension of these features varies from 16 (10 geometric + 6 chemical features as input) to 1 (binding score as output) throughout our network. Our data comes from the Protein Data Bank [7], with protein structures that are typically made up of A = 3K– 15K atoms and molecule sizes in the range 30 Å –300 Å (one ångström is equal to 10^−10^ m); we sample their surfaces at a resolution of 1 Å to work with N = 6K–15K points at a time.

**Figure 2:**
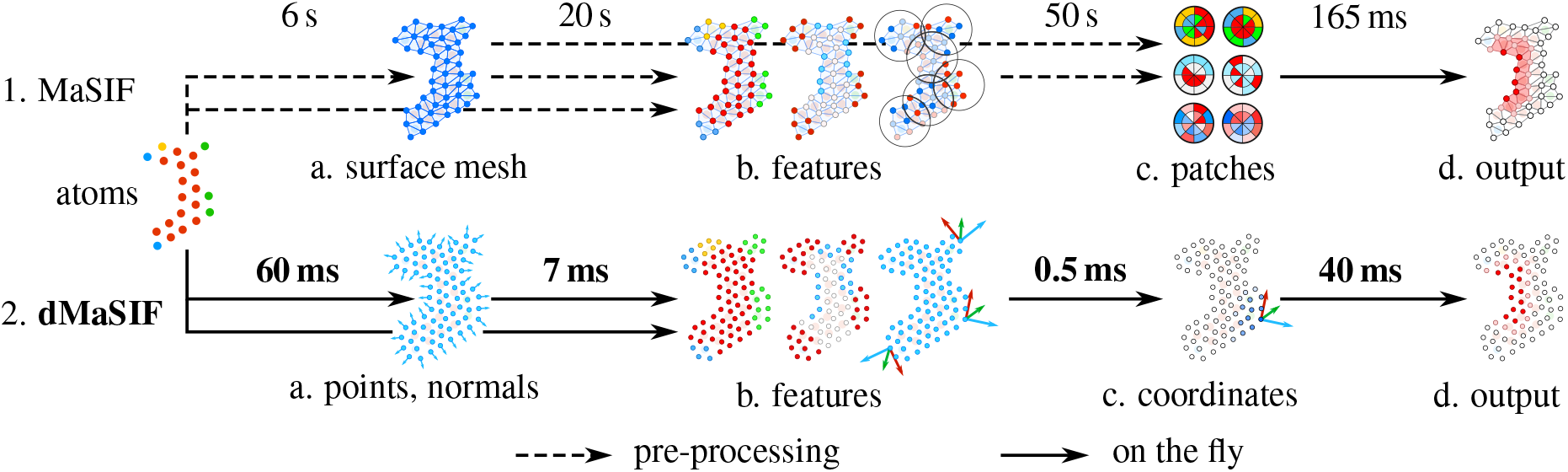
Both MaSIF and dMaSIF go through the same steps for interface prediction on protein surfaces. Starting from a raw atomic point cloud, we compute (a) a representation of the protein molecular surface, (b) geometric and chemical features, and (c) local coordinate systems; (d) a binding site is then predicted by a geometric convolutional neural network operating on (quasi-)geodesic patches on the protein surface. MaSIF precomputes steps (a)–(c), whereas we compute them on the fly 600 times faster. For every step, we display average run times per protein for inference on the site prediction task described in Section 4. Our method results in an accuracy level on par with MaSIF while alleviating the need for pre-calculations and providing significant speed-up for both inference and training.

We stress that unlike most other works for surface processing, our method *does not* rely on mesh structures, kNN graphs, or space partitioning of any kind. We compute exact interactions between all points of a protein surface efficiently using the recent KeOps library [13, 19] for PyTorch [30] that optimizes a wide range of computations on generalized distance matrices. ^1^

### 3.1. Surface generation

#### Fast sampling

The surface of a protein can be described as the level set of a smooth distance function or *meta ball* [9] (Figure 3a). To represent the six different atom types accurately, we associate an atomic radius *σ*_*k*_ to each atom **a**_*k*_ and define the smooth distance function:

**Figure 3:**
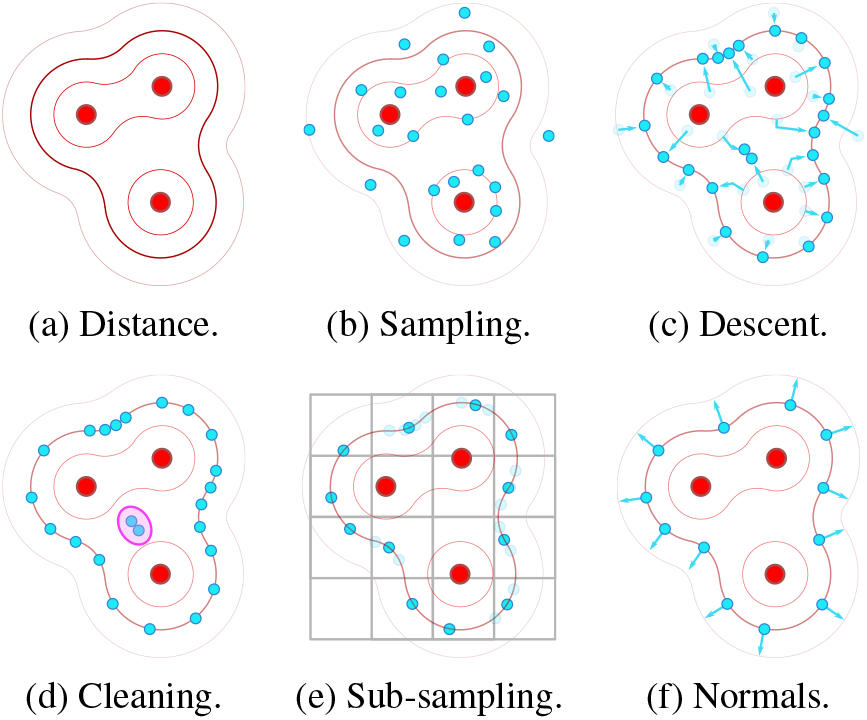
Sampling algorithm for protein surfaces. (a) Given the input protein (encoded as an atomic point cloud **a**_1_, *…*, **a**_A_, in red), its molecular surface is represented as a level set of the smooth distance function (1) to the atom centers. (b) To sample this surface, we first generate a point cloud **x**_1_, …, **x**_N=AB_ in the neighborhood of our protein (in blue): for every atom center, we draw B = 20 points from 𝒩 (*µ* = a_*k*_, *σ* = 10Å) and (c) let this random sample converge towards the target level set by gradient descent on (2) – we use 4 gradient steps with a learning rate of 1. (d) We then remove points trapped inside the protein: we keep a sample if the distance function at this location is close to our target value of *r* = 1.05 Å within a margin of 0.10 Å, and if making four consecutive steps of size 1 Å in the direction of the gradient of the distance function increases it by more than 0.5 Å. (e) We then put all points in cubic bins of side length 1 Å and keep one average sample per cell; this ensures that our sampling has uniform density. (f) Finally, the gradient of the distance function at location **x**_*i*_ is normalized to be used as a normal 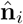.

**Figure 4:**
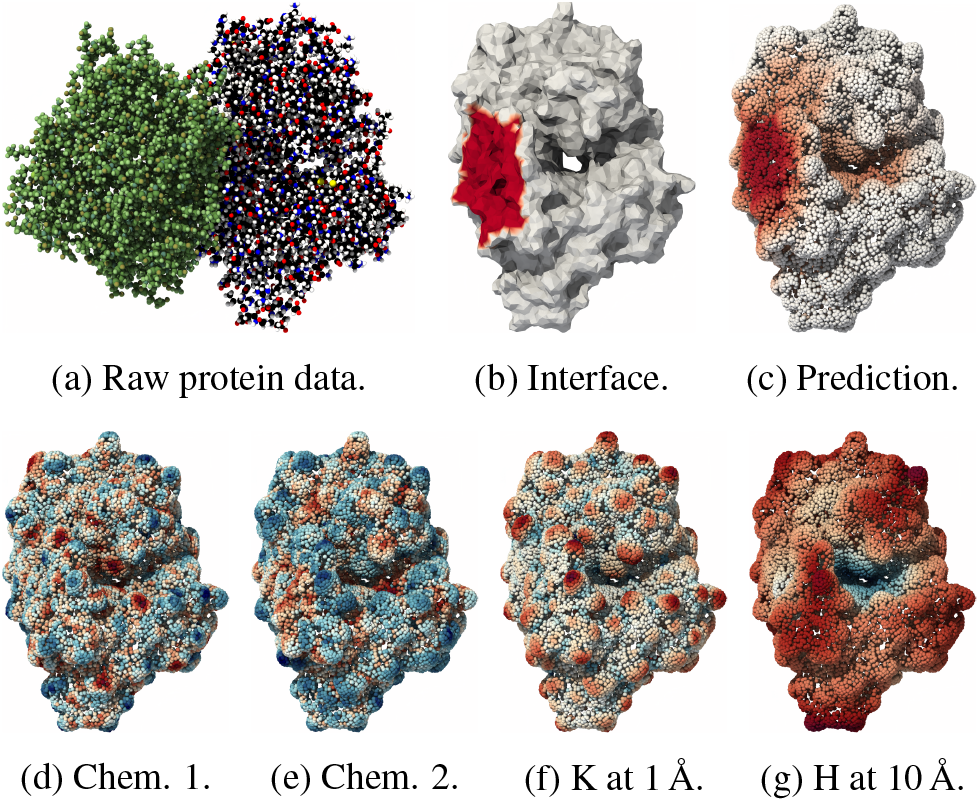
Illustration on the binding of the 1OJ7 pair. (a) The Protein Data Bank documents interactions between proteins 1OJ7_D (right) and 1OJ7_A (left, green). Can we learn to predict this 3D binding configuration from the unregistered structures of both proteins? (b) MaSIF tackles this problem as a surface segmentation problem. The binding site (red) is the ground truth signal that MaSIF tries to predict from precomputed chemical and geometric features, such as the electrostatic potential. It relies on mesh convolutions on the preprocessed molecular surface of the protein. (c) dMaSIF predicts the binding site without using any precomputed mesh structure or features. We perform all computations on an oriented point cloud, generated from the raw atom coordinates as in Figure 3. Data-driven chemical features (d-e) as well as Gaussian (f) and mean (g) curvatures at different scales are computed on the fly and given as inputs to a fast convolutional architecture that we describe in Figure 5. Rendering done with ParaView [5].

**Figure 5:**
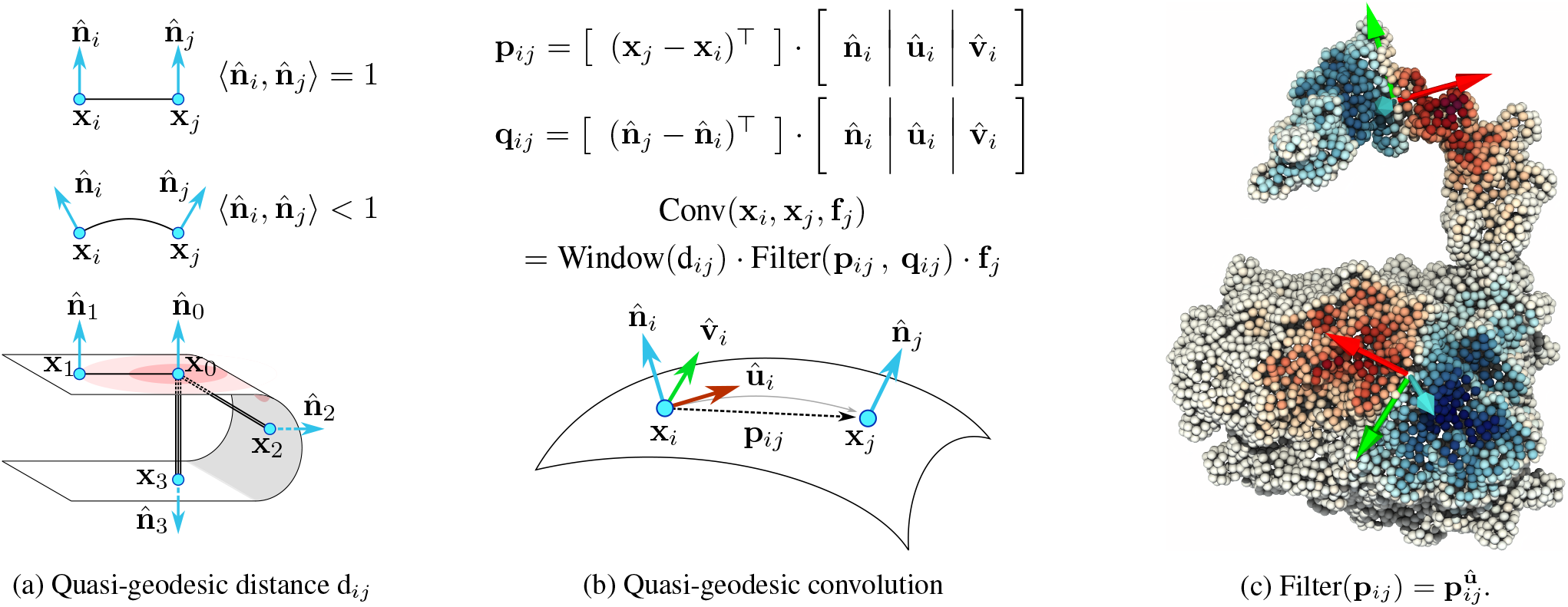
We use an approximation of the geodesic distance (5) to implement fast quasi-geodesic convolutions on oriented point clouds. (a) The weighted distance d_*ij*_ between points **x**_*i*_ and **x**_*j*_ is equal to | |**x**_*i*_ − **x**_*j*_ | | if the unit normal vectors 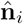 and 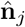 point towards the same direction, but is larger otherwise. In this example, the points **x**_1_, **x**_2_ and **x**_3_ lay at equal distance of the reference point **x**_0_ in ℝ^3^; but since the reference normal 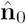 is aligned with 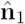, orthogonal to 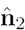 and opposite to 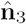, we have d_0,1_ = | |**x**_0_ − **x**_1_ | |*<* 2 d_0,1_ = d_0,2_ *<* 3 d_0,1_ = d_0,3_. (b) We leverage this behaviour to prevent information leakage “across the *volume*” of a protein. We combine a Gaussian window on the weighted distance d_*ij*_ with a parametric “Filter” to aggregate features **f**_*j*_ between neighbors on a protein *surface*. (c) Our formulae induce local coordinate systems that closely mimic the structure of genuine geodesic patches – defined here by a Gaussian window of deviation *σ* = 10 Å. On smooth surfaces, they enable the computation of “quasi-geodesic” convolutions at a much lower cost than mesh-based methods.

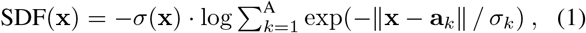

for any × ∈ ℝ^3^, with a stable log-sum-exp reduction and with 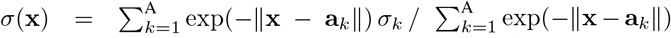 the average atom radius in a neighbourhood of point x.

As shown in Figure 3b, we sample the level set surface at radius *r* = 1.05 Å by minimizing the squared loss function:

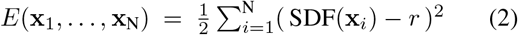

on a random Gaussian sample. KeOps allows us to implement this sampling strategy efficiently on batches of more than 100 proteins at a time (see Figure 6a).

**Figure 6:**
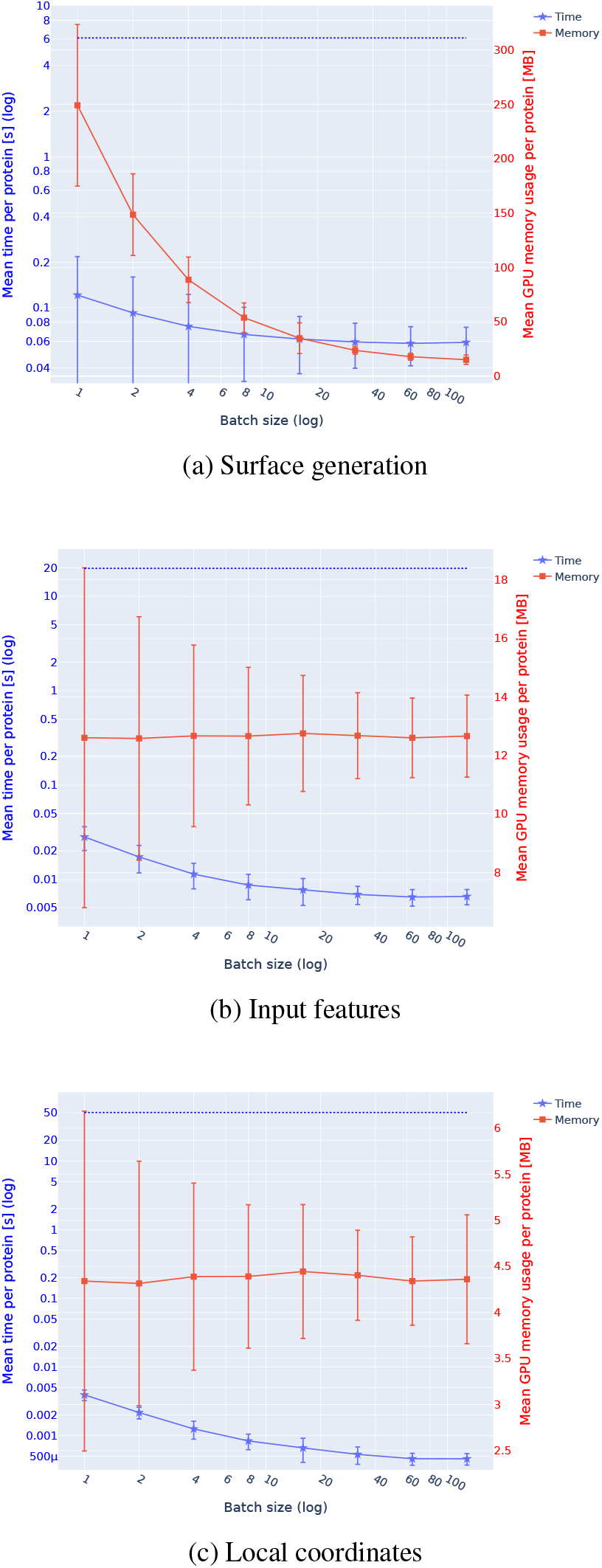
Computational cost of our “pre-processing” routines as functions of the batch size. We show the average time (blue curve and left axis, log scale) and memory (red curve, right axis, log scale) requirements of our method per protein, as a function of the number of proteins that are processed in parallel by our implementation. The dotted blue line shows the average time used by MaSIF to generate a surface mesh from the same atomic point cloud.

**Figure 7:**
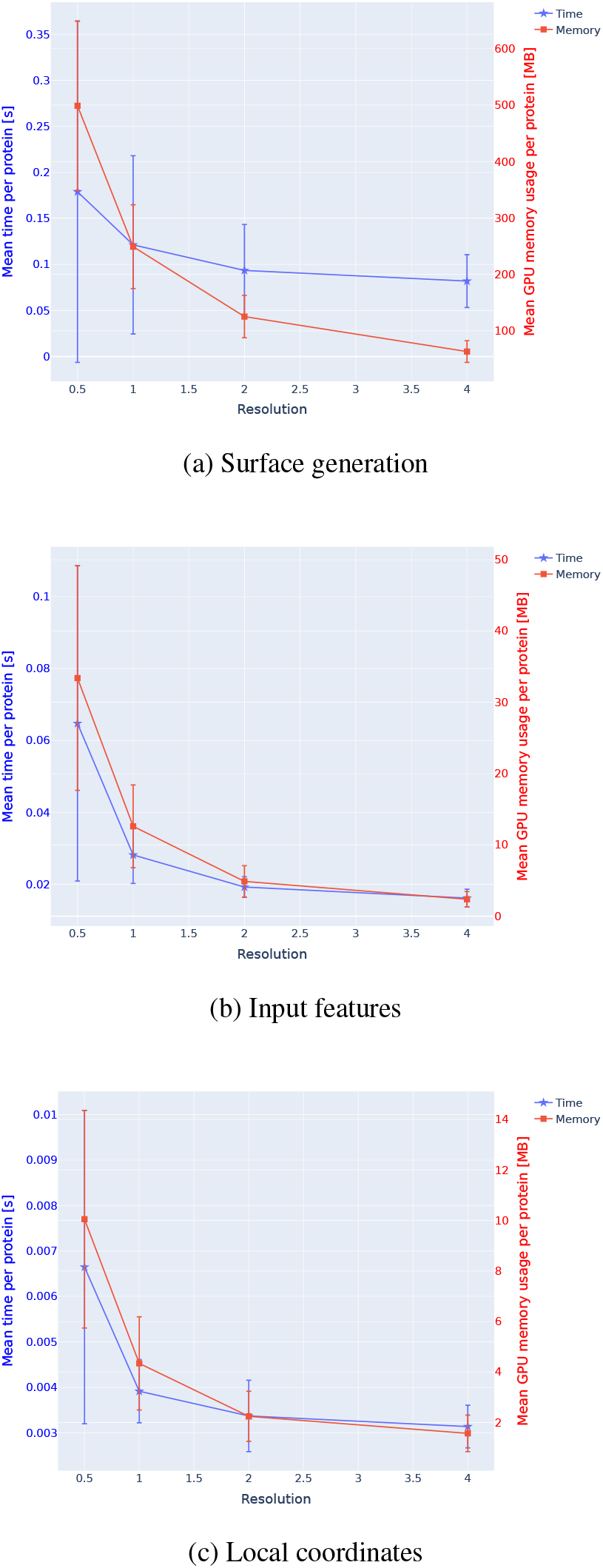
Computational cost of our “pre-processing” routines, as a function of the sampling resolution. We display the time (blue line and blue axis) and memory (red line and red axis) requirements of the pre-convolutional steps of our architecture as a function of the resolution of the generated point cloud. As expected, increasing the sampling density of our surface generation algorithm (i.e. using a lower resolution) results in longer processing times.

#### Descriptors

Point normals 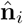 are computed using the gradient of the distance function (1). To estimate a local coordinate system 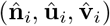, we first smooth this vector field using a Gaussian kernel with *σ* = 12 Å, i.e. use 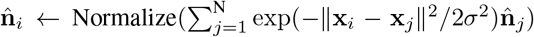. We then compute tangent vectors û_*i*_ and 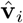 using the efficient formulae of [16]. Let 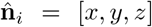 be a unit vector, *s* = sign(z), *a* = −1*/*(*s* + *z*) and *b* = *a × y*, then

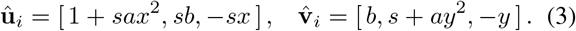

For each point **x**_*i*_, we then find the 16 nearest atom centers 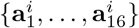 with types 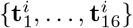 encoded as onehot vectors in ℝ^6^. We compute a vector of chemical features **f**_*i*_ in ℝ^6^ by applying a Multi-Layer Perceptron (MLP) to the vectors 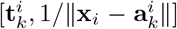 in ℝ^7^, performing a summation over the indices *k* = 1, *…*, 16 and applying a second MLP to the result. As illustrated in Figure 8, using simple MLPs with a single hidden layer of dimension 12 is enough to learn rich chemical features, such as the Poisson-Boltzmann electrostatic potential.

**Figure 8:**
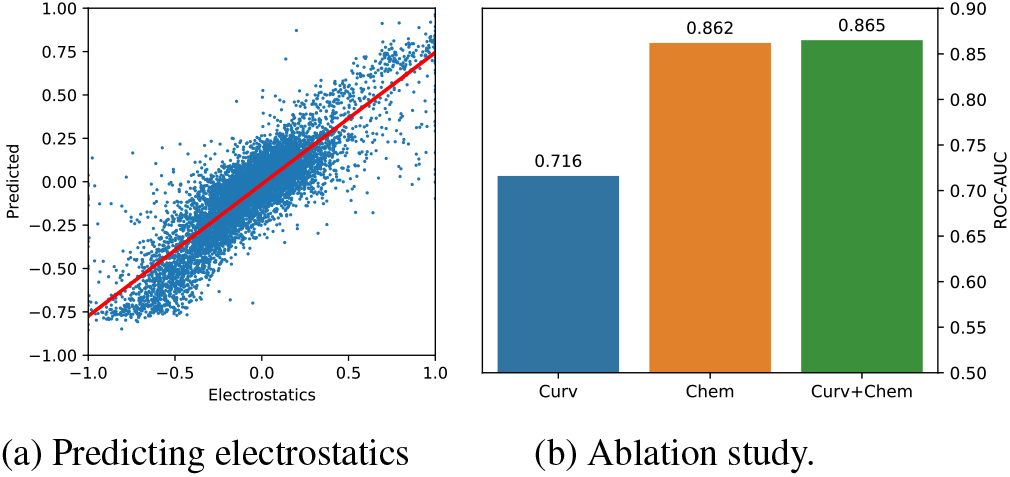
Our network can compute chemical properties of the protein surface from the underlying atomic point cloud. (a) Predicted Poisson-Boltzman electrostatic potential vs. the ground truth. Correlation cofactor r=0.83 and RMSE=0.16. (b) Ablation study showing how chemical and geometric features affect the performance in predicting interaction sites (ROC-AUC).

**Figure 9:**
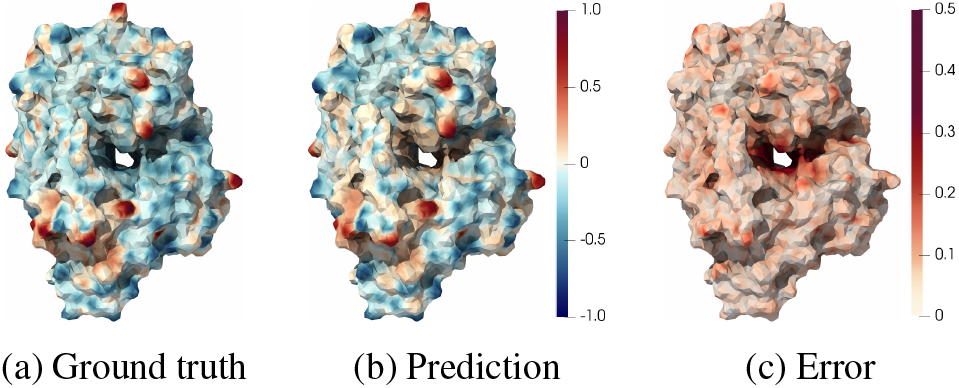
Comparison of groundtruth (a) and predicted (b) chemical properties of a protein. The error (c) is small, with RMSE=0.14. Most of the error is located inside the cavity.

### 3.2. Quasi-geodesic convolutions on point clouds

#### Convolutions on 3D shapes

To update the feature vectors **f**_*i*_ and progressively learn to predict the binding site of a protein, we rely on (quasi-)geodesic convolutions on the molecular surface. This allows us to ensure that our model is fully invariant to 3D rotations and translations, takes decisions according to *local* chemical and geometric properties of the surface, and is not influenced by atoms located deep inside the volume of a protein. These modelling hypotheses hold for many protein interaction problems and prevent our network from overfitting on the few thousands of protein pairs that are present in our dataset.

In practice, geometric convolutional networks combine pointwise operations of the form 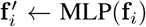 with local inter-point interactions of the form:

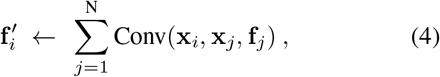

where **f**_*i*_ and 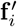 denote feature vectors associated to the point **x**_*i*_, and the “Conv” operator puts a trainable weight on the relationship between the points **x**_*i*_ and **x**_*j*_. The sum can possibly be replaced by a maximum or any other reduction operation.

#### Working with oriented point clouds

Numerous methods have been proposed to mimic surface operators with such convolution operators on meshes or point clouds – see Section 2 and especially [39, 25, 45, 40]. In this work, we leverage the reliable *normal vectors* produced by our sampling algorithm and the flexibility of the KeOps library to define a fast quasi-geodesic convolutional layer that works directly on oriented point clouds, *without any offline pre-computation* on the surface geometry.

As illustrated in Figure 5, we approximate the geodesic distance between two points **x**_*i*_ and **x**_*j*_ of a surface as:

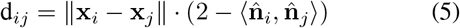

and localize our filters using a smooth Gaussian window of radius 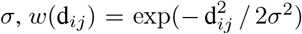. In the neighborhood of any point **x**_*i*_ of the protein surface, two 3D vectors then encode the relative position and orientation of neighbors **x**_*j*_ in the local coordinate system 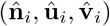.

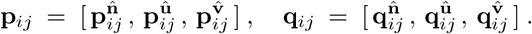

Different choices for the trainable “Filter” on these 3D vectors allow us to encode a wide range of operations. For the sake of computational efficiency, we focus on polynomial functions and MLPs instead of the popular Mixture-of-Gaussian filters [29], but note that this choice has little impact on the expressive power of our model.

#### Local orientation, curvatures

We must stress, however, that the pair of tangent vectors 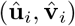 orthogonal to the normal 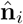 is defined up to a rotation of the tangent plane.

To work around this problem at a low computational cost, we follow [27] and orient the first tangent vector 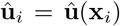 along the geometric gradient 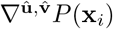 of a trainable potential *P* (**x**_*i*_) = *P*_*i*_ = MLP(**f**_*i*_), computed from the input features using a small MLP. We approximate its gradient using a derivative of Gaussian filter on the tangent plane, implemented as a quasi-geodesic convolution:

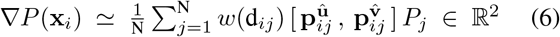

and then update the tangent basis 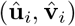 using standard trigonometric formulae.

Local curvatures are computed in a similar fashion [12]. We use quasi-geodesic convolutions with Gaussian windows of radii *σ* that range from 1 Å to 10 Å and quadratic filter functions to estimate the local covariances 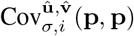 and 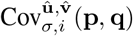 of the point positions and normals as 2 × 2 matrices in the tangent plane 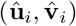. With *λ* = Å a small regularization parameter, the 2 2 shape operator at point **x**_*i*_ and scale *σ* is then approximated as 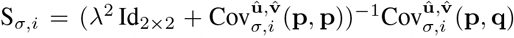, which allows us to define the Gaussian K_*σ,i*_ = det(S_*σ,i*_) and mean H_*σ,i*_ = trace(S_*σ,i*_) curvatures at scale *σ*.

#### Trainable convolutions

Finally, the main building block of our architecture is a quasi-geodesic convolution that relies on a trainable MLP to weigh features in a geodesic neighborhood of the local reference point **x**_*i*_. We turn a vector signal **f**_*i*_ ∈ ℝ^F^ into a vector signal 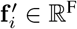 with:

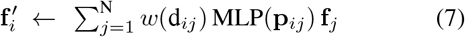

where MLP is a neural network with 3 input units, H = 8 hidden units, ReLU non-linearity and F = 16 outputs.

### 3.3. End-to-end convolutional architecture

#### Overview

We chain together the operations introduced in the previous sections to create a fully differentiable pipeline for deep learning on protein surfaces, illustrated in Figure 2. As a brief summary:

1. We sample surface points and normals as in Figure 3.
2. We use the normals 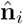 to compute mean and Gaussian curvatures at 5 scales *σ* ranging from 1 Å to 10 Å.
3. We compute chemical features on the protein surface as described in Section 3.1. Atom types and inverse distances to surface points are passed through a small MLP with 6 hidden units, ReLU non-linearity and batch normalization [24]. Contributions from the 16 nearest atoms to a surface point **x**_*i*_ are summed together, followed by a linear transformation to create a vector of 6 scalar features.
4. We concatenate these chemical features to the 5 + 5 mean and Gaussian curvatures to create a full feature vector of size 16.
5. We apply a small MLP on this vector to predict orientation scores *P*_*i*_ for each surface point. We then orient the local coordinates 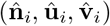 according to (6).
6. We apply successive trainable convolutions (7), MLPs and batch normalizations on the feature vectors **f**_*i*_. The numbers of layers, the radii of the Gaussian windows and the number of units for the MLPs are task-dependent and detailed in the Supplementary Material.
7. As a final step for site identification, we apply an MLP to the output of the convolutions to produce the final site/non-site binary output. For interaction prediction, we compute dot products between the feature vectors of both proteins to use them as interaction scores between pairs of points.

#### Asymmetry between binding partners

When trying to predict binding interactions for protein pairs, we process both interacting proteins identically up to the convolutional step. We then introduce some asymmetry by passing each one of the two binding partners through a separate convolutional network. This allows the network to find *complementary* (instead of similar) regions on both surfaces, such as convex bulges and concave pockets. We note that MaSIF encoded such an asymmetry by inverting the sign of the pre-computed features on one of the two surfaces.

## 4. Experimental Evaluation

### Benchmarks

We test our method on two tasks introduced in [20]. The tasks come from the field of structural bioinformatics and deal with predicting how proteins interact with each other.

#### Binding site identification

we try to classify the surface of a given protein into interaction sites and non-interaction sites. Interaction sites are surface patches that are more likely to mediate interactions with other proteins: understanding their properties is a key problem for drug design and the study of protein interaction networks. The identification of the interaction site is unaware of the binding partner.

#### Interaction prediction

we take as inputs two surface patches, one from each protein involved in a complex, and predict if these locations are likely to come into close contact in the protein complex. This task is key to prediction tasks like protein docking, i.e. predicting the orientation of two proteins in a complex.

### Dataset

The dataset comprises protein complexes gathered from the Protein Data Bank (PDB) [7]. We use the training / testing split of [20], which is based on sequence and structural similarity and was assembled to minimize the similarity between structures of the interfaces in the training and testing set. For site identification, the training and test sets include 2958 and 356 proteins, respectively; 10% of the training set is reserved for validation. For interaction prediction, the training and test sets include 4614 and 912 protein complexes, respectively, with 10% of the training set used for validation.

The average number of points used to represent a protein surface is N = 11549 ± 1853 for our generated point clouds, compared to 6321 ± 1028 points for MaSIF.^2^ Proteins are randomly rotated and centered to ensure that methods which rely on atomic point coordinates do not overfit on their spatial locations.

### Baselines

Our main baselines are the MaSIF-site and MaSIF-search models [20]. For the MaSIF baselines, we use the pre-trained models and precomputed surface meshes and input features provided by the authors. Additionally, in order to show the benefits of our convolutional layer, we benchmark it against PointNet++ [34] and Dynamic Graph CNN (DGCNN) [43], two popular state-of-the-art convolutional layers for point clouds.

### Implementation

We implement our architectures with PyTorch [30] and use KeOps [19] for fast geometric computations. For data processing and batching, we use Py-

Torch Geometric [18]. For the PointNet++ and DGCNN baselines, we use PyTorch Geometric implementations – but rely on KeOps symbolic matrices to accelerate the construction of kNN graphs and thus guarantee a fair comparison. For the MaSIF baselines, we use the reference implementation of [20].^3^ All models are trained on either a single NVIDIA GeForce RTX 2080 Ti GPU or a single Tesla V100. Run times and memory consumption are measured on a single Tesla V100.

### 4.1. Surface and input feature generation

#### Precomputation

A key drawback of MaSIF is its reliance on the heavy precomputation of surface meshes and input features. These computations take a significant amount of time and generate large files that must be stored on disk. For reference, the pre-processed files used to train the MaSIF networks weigh more than 1TB. In sharp contrast, our method does not rely on any such pre-computation. Table 1 compares corresponding run times for both pipelines: our method is three orders of magnitude faster than MaSIF for these geometric computations.

**Table 1:**
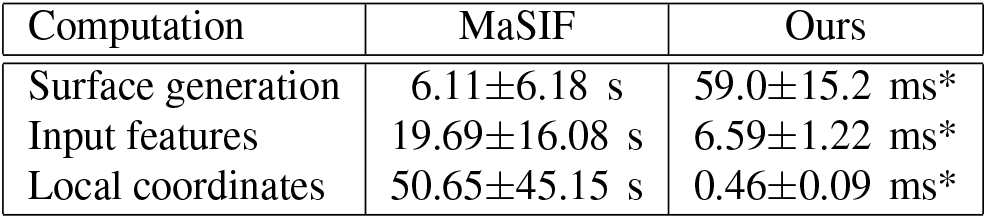
Average “pre-processing” time per protein. Our method is about 1000 times faster than MaSIF and allows these computations to be performed on the fly, as opposed to the offline precomputations of MaSIF. *With batches of 128 proteins at a time.

| Computation | MaSIF | Ours |
| --- | --- | --- |
| Surface generation | 6.11±6.18 s | 59.0±15.2 ms* |
| Input features | 19.69±16.08 s | 6.59±1.22 ms* |
| Local coordinates | 50.65±45.15 s | 0.46±0.09 ms* |

#### Scalability

Our surface generation algorithm scales beneficially with an increasing batch size. In Figure 6, we show that the running time and memory requirement per protein of our method both decrease significantly when processing dozens of proteins at time the batch size. This is a consequence of the increased usage of the GPU cores and the smaller influence of fixed PyTorch and KeOps overheads.

Moreover, our method of surface generation makes it easy to experiment with different point cloud resolutions. Different tasks could benefit from higher or lower resolution and tuning it as a hyperparameter could have significant effects on performance. We show the effects of resolution on time an memory requirements in Figure 7.

#### Quality of learned chemical features

Another notable drawback of MaSIF is its reliance on ‘handcrafted’ geometric and chemical features (Poisson-Boltzmann electrostatic potential, hydrogen bond potential and hydropathy) that must be precomputed and provided as input to the neural network. In contrast, we do not use any handcrafted descriptors and learn problem-specific features directly from the underlying atomic point cloud, provided as the sole input of our method. We argue that this information alone is sufficient to compute an informative chemical and geometric description of the protein surface. To support this statement, we show in Figure 8 the results of an experiment where our chemical feature extractor is used to regress the Poisson-Boltzmann electrostatic potential on surface points. The quality of our predicition suggests that our data-driven chemical features are of similar quality to the descriptors used by MaSIF – or better.

We also note the results of an ablation study for chemical and geometric features, depicted in Figure 8. They suggest that the concatenation of geometric curvatures to the vector of learned chemical features does not significantly improve the performance of the network for the site prediction task: we will investigate this point in future works.

### 4.2. Performance

#### Binding site identification

Results for the identification of binding sites are summarized in Figures 10–12, which depict ROC curves and tradeoffs between accuracy, time and memory. We evaluate multiple versions of our architecture with varying numbers of convolution layers (1 vs 3) and patch sizes (5, 9, or 15Å). For comparison, we also show results when our convolutions are replaced by DGCNN and PointNet++ architectures, all other things being equal.

**Figure 10:**
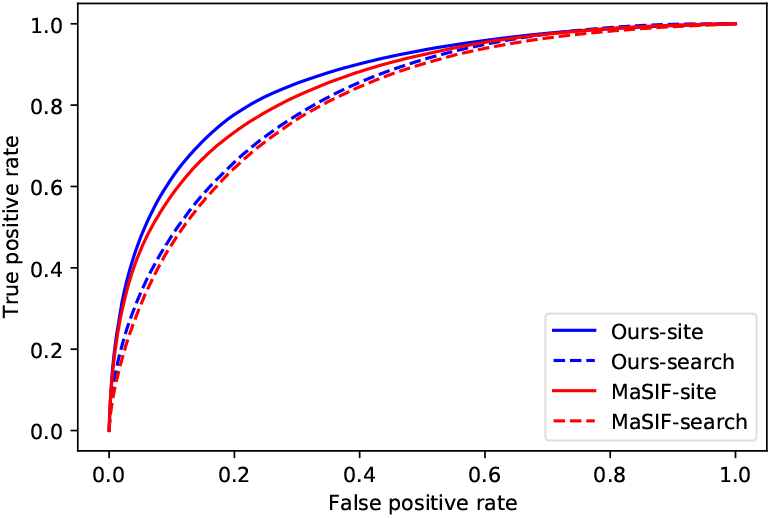
ROC curves comparing the performance of our method (blue) and MaSIF (red) on the task of binding site identification (solid curves) and search of binding partners (dashed). Our approach performs on par with MaSIF, achieving ROC-AUC of 0.87 (vs. 0.85) in site identification, and 0.82 (vs. 0.81) in identifying binding partners.

**Figure 11:**
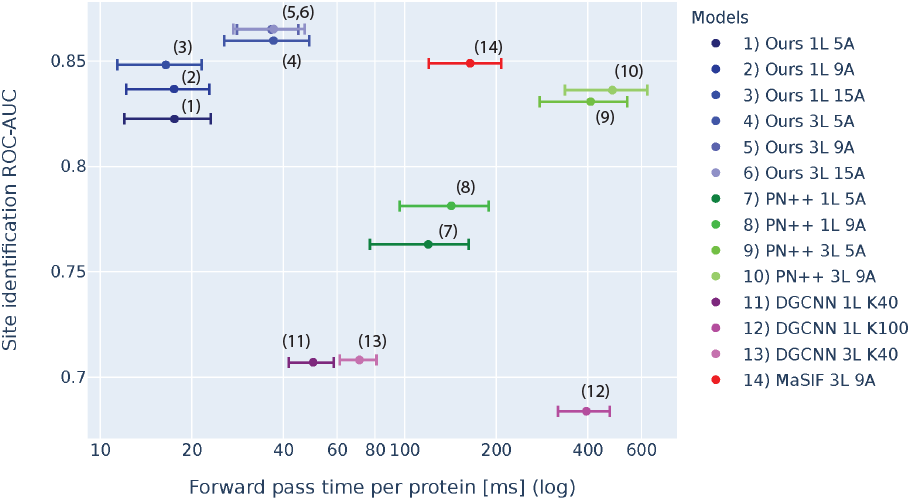
Accuracy (site identification ROC-AUC) vs. Run time (forward pass/protein in ms) of different architectures.

**Figure 12:**
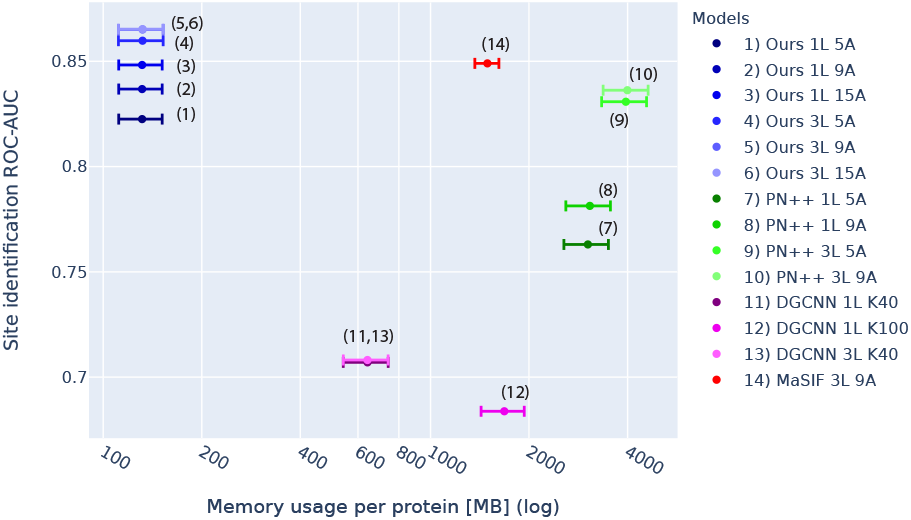
Accuracy (site identification ROC-AUC) vs. Memory footprint (MB/protein) of different architectures.

A first remark is that if we use a single convolution layer with a Gaussian window of deviation *σ* = 15 Å, our method matches the best accuracy of 0.85 ROC-AUC produced by MaSIF – with 3 successive convolutional layers on patches of radius 9 Å. In this configuration, our network runs 10 times faster than MaSIF with an average time in the forward pass of 16 ms vs. 164 ms per protein. At the price of a modest increase of the model complexity (three convolution layers, and 36 ms on average per protein), we outperform MaSIF with a 0.87 ROC-AUC, detailed in Figure 10 (solid curves). Most remarkably, our models all have a small memory footprint (132 MB/protein), which is 11 times less than an equivalent MaSIF network (1492 MB/protein), 13 times less than DGCNN (1,681 MB/protein) and 30 times less than PointNet++ (3,995 MB/protein).

#### Interaction prediction

With a single convolutional layer architecture similar to that of MaSIF-search we reach a slightly higher performance of 0.82 vs. 0.81, as illustrated in Figure 10 (dashed). We remark that MaSIF-search reaches this level of accuracy using high dimensional feature vectors with 80 dimensions compared to our 16: understanding the influence of the number of convolutional “channels” on the performances of our network for different tasks will be an important direction for future works.

Note that MaSIF-search also relies on larger patches than MaSIF-site (12 Å vs. 9 Å), which causes a significant increase of run times to 727 ± 403 ms. On the other hand, our lightweight method runs in 17.5±6.7 ms and is over 40 times faster at inference time.

## 5. Conclusion

We have introduced a new geometric architecture for deep learning on protein surfaces, enabling the prediction of their interaction properties. Our method is an order of magnitude faster and more memory efficient than previous approaches, making it suitable for the analysis of large-scale datasets of protein structures: this opens the door to the analysis of entire protein-protein interaction networks in living organisms, comprising over 10K proteins.

The fact that our pipeline works on raw atomic coordinates and is fully differentiable makes it amenable to *generative* tasks, with the possibility of performing a true end-to-end design of new proteins for diverse biological functions, namely in terms of the design of binders for specific targets. This opens fascinating perspectives in drug design, including biologics for targeting disease relevant targets (e.g. cancer therapy, antiviral) that display flat interaction surfaces and are impossible to target with small molecules.

More broadly, we believe that our new algorithmic and architectural ideas for deep learning on 3D shapes through fast on-the-fly computations on point clouds will be of general interest to computer vision and graphics experts. Conversely, we hope that our work will draw the attention of this community to some of the most important and promising problems in structural biology and protein science.

## Footnotes

1 The size 5K–20K and dimension 3 of our point clouds appear to be a sweetspot for KeOps in ‘bruteforce mode’, thanks to *contiguous* operations that stream much better on GPUs than the *scattered* memory accesses of graph-based and hierarchical methods.

2 This smaller sampling size of MaSIF stems from the large time and memory requirements of this method, which prohibits the use of finer meshes.

3 Since MaSIF is implemented in TensorFlow [1], small discrepancies in measurements of memory consumption and running times are possible.

