## Supplementary for "Fast end-to-end learning on protein surfaces"

### Supplementary material

#### 1. Description of network architectures

A high level description of our networks for both site identification and interaction prediction can be found in Figs. 1 and 2 respectively. In these diagrams, "FC(I,O)" denotes a fully connected (linear) layer with I input channels and O output channels; "LR" denotes a Leaky ReLU activation function with a negative slope of 0.2; "BN" denotes a batch normalization layer. Red, blue and green blocks denote atom properties, surface descriptors and feature vectors, respectively.

We estimate chemical features on the generated surface points using the architecture described in Fig. 4. This module takes as inputs the atom coordinates and types, along with the surface point coordinates. For each point on the surface, the network finds the 16 nearest atoms and assigns a 6-dimensional chemical feature based on the atom types and their distances to the point. As detailed in Fig. 3, we concatenate these chemical features to a 10-dimensional vector of geometrical features, which approximate the mean and Gaussian curvatures at different scales.

We then pass these input feature vectors through a sequence of convolutional layers (Fig. 5). As discussed in Section 3 of the paper, we first use the surface normals $\hat{\mathbf{n}}_i$ to build local tangent coordinate systems and orient the unit tangent vectors $\hat{\mathbf{u}}_i$, $\hat{\mathbf{v}}_i$ according to the gradient of an orientation score $P_i$. Finally, we use this complete description of the surface geometry to establish quasi-geodesic convolutional windows and progressively update our feature vectors.

The DGCNN and PointNet++ baselines replace the "convolutional" block of our architecture with standard alternatives provided by PyTorch Geometric. We keep the same numbers of channels as for our method (8 for the site prediction task, 16 for the search predicition task) and benchmark runs with several interaction radii and number of K-nearest neighbors.

#### 2. Description of the training process

We filter the datasets according to the criteria described in [1]. To be considered in our benchmarks, each protein must have at least 30 interface points and the interface has to cover less than 75% of the total surface area.

| Parameter | Site | Search |
|---|---|---|
| Optimizer | AMSGrad | AMSGrad |
| Learning rate | $3 \times 10^{-4}$ | $3 \times 10^{-4}$ |
| Epochs | 50 | 100 |
| Descriptor dimensionality | 8 | 16 |
| Early stopping | Yes | Yes |

Table 1: Hyperparameters for our training loops.

**Binding site identification.** We detail our hyperparameters in Table 1. Surfaces are generated in batches, but predictions are only performed on single proteins at a time. From each protein, 16 positives and 16 negatives locations are randomly sampled and the loss function is computed on these points. We found that this process stabilized the training process and improved generalization. Labels are mapped from precomputed MaSIF meshes by finding the nearest neighbours. Furthermore, if a point is further than 2.0Å away from any precomputed mesh point, it is labeled as non-interface. The loss is computed as the binary cross entropy between the labels and the predictions.

**Interaction prediction.** Surface generation and prediction are performed in the same way as for binding site identification. However, as detailed at the end of Section 3.3 in the paper, each binding partner is passed through a separate convolutional network. The prediction scores are then computed by taking the inner product between the convolutional embeddings of the two proteins. Pairs of points are labeled as interacting if they are less than 1Å from each other. From each protein, 16 positives and 16 negatives were randomly sampled. The loss was computed as the binary cross entropy.

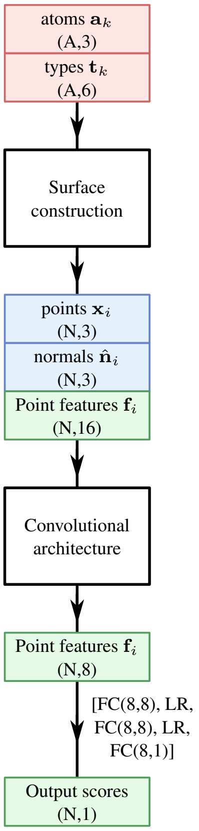


Figure 1: Overview of our architecture for the site prediction task, that we handle as a binary classification problem of the surface points. The "surface construction" block is detailed in Figure 3, while the "convolutional architecture" is detailed in Figure 5.

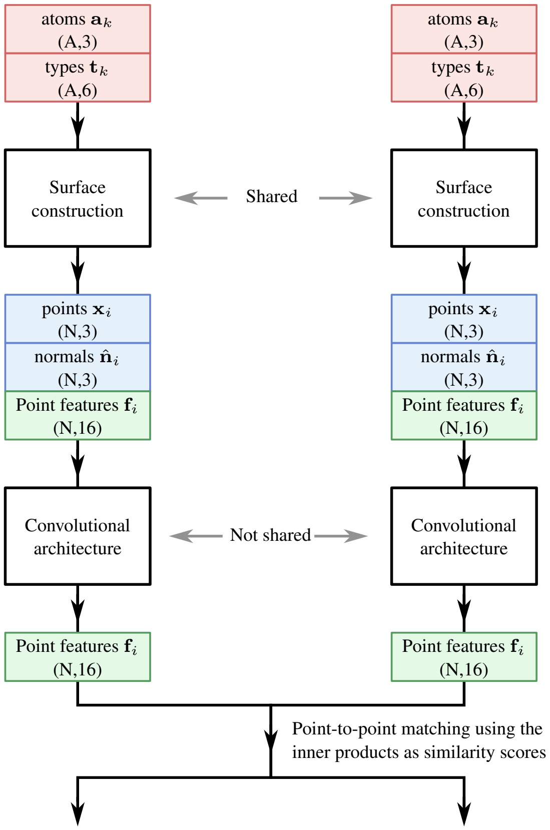


Figure 2: Overview of our architecture for the search prediction task. The "surface construction" block is detailed in Figure 3, while the "convolutional architecture" is detailed in Figure 5.

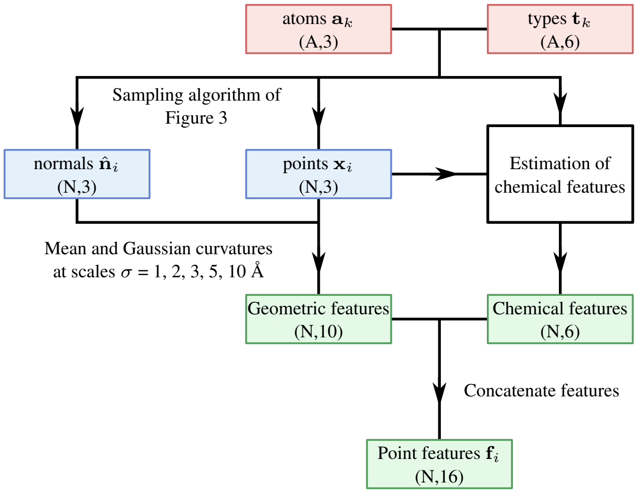


Figure 3: Construction of a surface representation, detailed in Section 3.1 of the paper. The “chemical features” block is detailed in Figure 4.

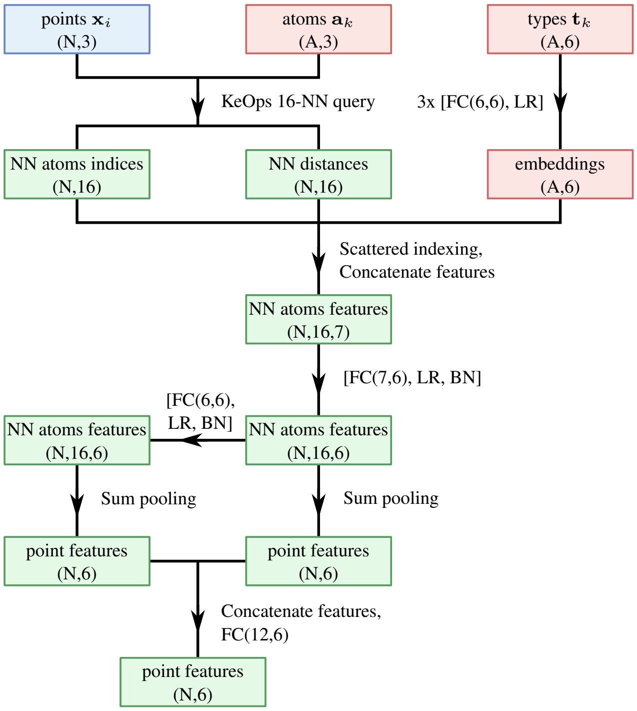


Figure 4: Estimation of chemical features from the raw atom types and coordinates.

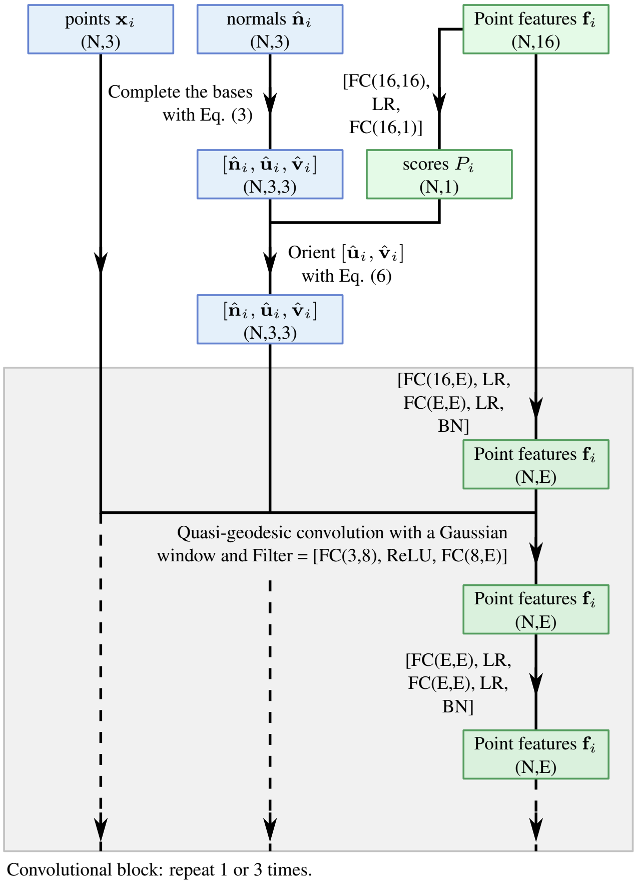


Figure 5: Convolutional architecture, with E convolutional “channels” (we use E=8 for the site prediction task and E=16 for the search prediction task). Our architecture for the search prediction task has an additional skip connection between the inputs and outputs. As detailed in Section 3.2, our network first estimates local coordinate systems $[\hat{\mathbf{n}}_i, \hat{\mathbf{u}}_i, \hat{\mathbf{v}}_i]$ attached to the points $\mathbf{x}_i$ of a protein surface. We then rely on a fast approximation of the geodesic distance to define quasi-geodesic convolutions and let our feature vectors $\mathbf{f}_i$ interact on the protein surface.

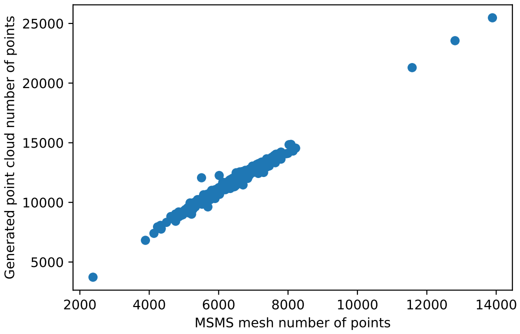


(a) Number of points.

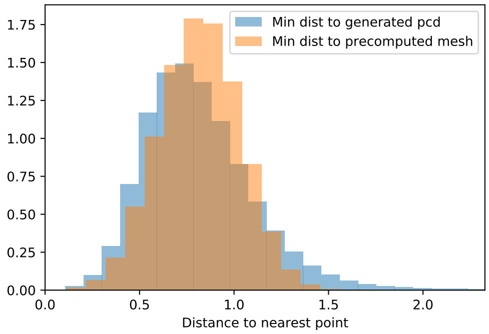


(b) Distances to closest points.

Figure 6: Quality control for our surface generation algorithm. (a) Number of points generated per protein by our method, as a function of number of points in the precomputed mesh used by MaSIF. As expected, we observe a nearly perfect linear correlation. (b) For each point generated by our method, we display in orange the distance to the closest point on the precomputed mesh. Conversely, we display in blue the histogram of distances to the closest generated point, for points on the MaSIF "ground truth" mesh. We noticed that the blue curve showed a very long tail (not visible on this figure). This comes from an artifact in the surface generation algorithm of MaSIF, which cuts out parts of proteins that have missing densities. We solved this discrepancy by removing these points from our dataset as well, and only display point-to-point distances in the 99th percentile – i.e. we treat the largest 1% distances as outliers, not displayed here.

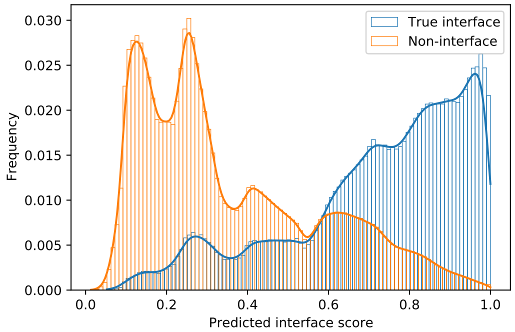


Figure 7: Additional display for the site prediction task. We display the distributions of predicted interface scores for both true interface points (blue) and non-interface points (orange). The separation is clear, resulting in a ROC-AUC of 0.87 in Figure 8 of the paper.
